# Interferon-gamma and TNF-alpha synergistically enhance the immunomodulatory capacity of Endometrial-Derived Mesenchymal Stromal Cell secretomes by differential microRNA and extracellular vesicle release

**DOI:** 10.1101/2021.06.10.447490

**Authors:** María de los Ángeles de Pedro, Federica Marinaro, Esther López, María Pulido, Francisco Miguel Sánchez-Margallo, Verónica Álvarez, Javier G Casado

## Abstract

Endometrial Mesenchymal Stromal Cells (endMSCs) can be easily isolated from menstrual blood by plastic adherence. These cells have a potent pro-angiogenic and immunomodulatory capacity, and their therapeutic effect is mediated by paracrine mechanisms where secretome have a key role. In this paper, we aimed to evaluate different priming conditions in endMSCs using pro-inflammatory cytokines and Toll-Like Receptor ligands. Our *in vitro* results revealed a synergistic and additive effect of IFNγ and TNFα on endMSCs. The combination of these pro-inflammatory cytokines significantly increased the release of Indoleamine 2,3-dioxygenase (IDO1) in endMSCs. Additionally, this study was focused on the phenotype of IFNγ/TNFα-primed endMSCs (endMSCs*). Here we found that immune system-related molecules such as CD49d, CD49e, CD54, CD56, CD58, CD63, CD126, CD152, or CD274 were significantly altered in endMSCs* when compared to control cells. Afterward, our study was completed with the characterization of released miRNAs by Next Generation Sequencing (NGS). Briefly, our system biology approaches demonstrated that endMSCs* showed an increased release of 25 miRNAs whose target genes were involved in immune response and inflammation. Finally, the cellular and molecular characterization was completed with *in vitro* functional assays.

In summary, the relevance of our results lies in the therapeutic potential of endMSCs*. The differences in cell surface molecules involved in migration, adhesion and immunogenicity, allowed us to hypothesize that endMSCs* may have an optimal homing and migration capacity towards inflammatory lesions. Secondly, the analysis of miRNAs, target genes and the subsequent lymphocyte activation assays demonstrated that IFNγ/TNFα-primed secretome may exert a potent effect on the regulation of adverse inflammatory reactions.

## Introduction

Mesenchymal stromal cells (MSCs) are multipotent fibroblast-like plastic-adherent cells (1) that can be isolated from various pre- and post-natal tissues (2), being the endometrium among them. Endometrial-derived MSCs (hereinafter referred to as endMSCs) can be isolated from biopsies containing the endometrial *stratum functionalis* and *stratum basalis*, or from menstrual fluid (3). These cells can be further purified by plastic adherence and *in vitro* expanded using standard culture conditions (4). The scientific literature has also described different stem/progenitor cells in the three tissue layers of endometrium (5–7) and how to isolate them using different procedures (8–10). Isolation from menstrual blood is a non-invasive and reproducible method (5) that allows the retrieval of MSCs fulfilling the minimal criteria proposed by the International Society of Cellular Therapy. Firstly, they have plastic adherence in tissue culture. Secondly, they display a CD105+, CD73+, CD90+, CD45-, CD34-, CD14-, and HLA-DR-phenotype. Third, they possess an *in vitro* differentiation capacity towards osteoblasts, adipocytes and chondroblasts (11).

MSCs have well-known immunomodulatory and regenerative properties (12), being used in preclinical and clinical trials (13). Moreover, these endMSCs have also demonstrated an immunosuppressive activity (14–16), as wells as anti-apoptotic and pro-angiogenic capacities (17) which are mediated by paracrine factors. In this paracrine activity, secretome is necessary for intercellular communication and comprise extracellular vesicles (EVs), exosomes, proteins, nucleic acids, and lipids with therapeutic potential.

Many strategies to improve the regenerative effects and efficacy of secretome components have been recently proposed and reviewed (18,19). Basically, the main idea is to optimize *in vitro* culture conditions and provide stimuli (the so-called “priming” or “licensing”) to use MSCs as “cell factories”. In this process, the “manufactured products” are therapeutically bioactive vesicles. Nowadays, most of the priming strategies have been optimized using MSCs derived from bone marrow, from umbilical cord, or adipose tissue (19). These priming strategies have been evaluated using different cytokines, such as IL17A, IL1β, FGF2, TNFα, and IFNγ (20–24); different molecules, such as LL37, Lipopolysaccharide (LPS), Polyinosinic: polycytidylic acid (Poly (I:C)), curcumin, oxytocin, and melatonin (25–28); different chemical agents such as 5-aza-2′-deoxycytidine, valproic acid, and sphingosine-1-phosphate (29–31); and different hypoxia conditions (32–36). Additionally, all these priming strategies have been combined and optimized using different concentrations of the priming agent. For example, LPS has been used in combination with TNFα (21) or Poly(I:C) in MSCs with effects on cytokine secretion (37).

There are plenty of studies focused on the MSC-priming strategy with very different purposes: induction of angiogenesis, regeneration, or cell viability. In this study, we have evaluated different priming strategies to enhance the immunomodulatory capacity of secretomes from endMSCs. Moreover, the immunomodulatory effect of their secreted vesicles has already been demonstrated by our group (38), and additional studies allowed us to understand, at least in part, the complexity of the molecular networks and their effects on target cells (39).

There are several publications focused on the analysis of IFNγ- and TNFα-primed cells. In two of them, authors demonstrated an upregulation of indoleamine 2,3-dioxygenase (IDO1) (40), which is mediated by chromatin remodeling at the *IDO1* promoter (41). Moreover, an inhibition of complement activation by factor H has been described (42). It is important to note that these publications analyzed the effect of priming upon cells, but not upon the secreted vesicles.

On the other hand, priming strategies using IFNγ and TNFα have also been widely studied for secretome based therapies. Here we summarize some studies focused on the analysis of secretomes by IFNγ/TNFα-primed MSCs. In the case of IFNγ, EVs from IFNγ-primed MSCs have been found to enhance macrophage bacterial phagocytosis (43) and membrane particles obtained from IFNγ-primed MSCs can induce an increase of anti-inflammatory PD-L1 monocytes (44). In the case of TNFα, EVs from TNFα-primed MSCs showed a therapeutic effect in urethral fibrosis (45), an improvement in the proliferation and differentiation of osteoblasts (46), as well as neuroprotective effects (47). In the case of priming strategies using the combinations of IFNγ/TNFα, *in vitro* studies have revealed that IFNγ/TNFα-primed MSCs produced EVs with a heightened immunomodulatory potential due to prostaglandin E2 and cyclooxygenase 2 pathway alteration (48). Priming MSCs with IFNγ/TNFα also shifted macrophages polarization from M1 to M2 by miRNAs (49) and resulted in immunomodulatory EVs which induced M2 differentiation and enhancement of Tregs (50). All these studies suggest that IFNγ and TNFα have an enormous potentiality to act as priming agents for MSCs.

According to our previous studies (38,39) and considering all preceding findings in different MSCs, the aim of this study was to evaluate different priming strategies in endMSCs. Our results firstly revealed a synergistic and additive effect of IFNγ and TNFα that significantly triggered the release of IDO1. Secondly, these cytokines significantly contributed to phenotypic changes in molecules involved in migration, adhesion, and immunogenicity. Third, IFNγ/TNFα-primed endMSCs (endMSCs*) deliver miRNAs which target inflammation-related genes. Finally, *in vitro* functional assays have demonstrated the immunomodulatory capacity of these vesicles.

To our knowledge, this is the first report focused on the optimization of priming strategies in endMSCs. Our results have demonstrated that IFNγ/TNFα priming have a profound impact on the immunomodulatory potential of released vesicles and miRNAs profile of these secretomes, as well as in the migration/adhesion capacity of these cells. Altogether, these results may have clinical implications for the therapeutic use of secretomes in inflammatory mediated-diseases.

## Material and Methods

### Isolation, culture, and characterization of human endMSCs

A summary of all the experimental procedures is shown in Figure 1. Written informed consent was obtained from human menstrual blood donors and approved by the Ethics Committee of the Jesús Usón Minimally Invasive Surgery Center. Human menstrual blood samples were collected by three healthy pre-menopausal women and endMSCs were isolated under sterile conditions according to a previously described protocol (38). Following a 1:2 dilution and homogenization of the menstrual blood in phosphate buffered saline (PBS), a first centrifugation at 450 × g for 10 min was carried out. The pelleted cells were recovered and seeded in Dulbecco’s Modified Eagle’s medium (DMEM) containing 10% fetal bovine serum (FBS) (Gibco, Thermo Fisher Scientific, Bremen, Germany), 1% penicillin/streptomycin, and 1% glutamine, at 37 °C and 5% CO_2_. The adherent endMSCs were cultured to 80% confluency, onto tissue culture flasks, while the non-adherent cells were discarded after 24 h. The cell culture medium was replaced every 3 or 4 days. The cells were detached from tissue culture flasks using PBS containing 0.25% trypsin (Lonza, Gaithersburg, MD, USA). The isolated endMSCs were characterized by flow cytometry and differentiation assay, as in our previous studies (38,39,51).

**Figure 1.**
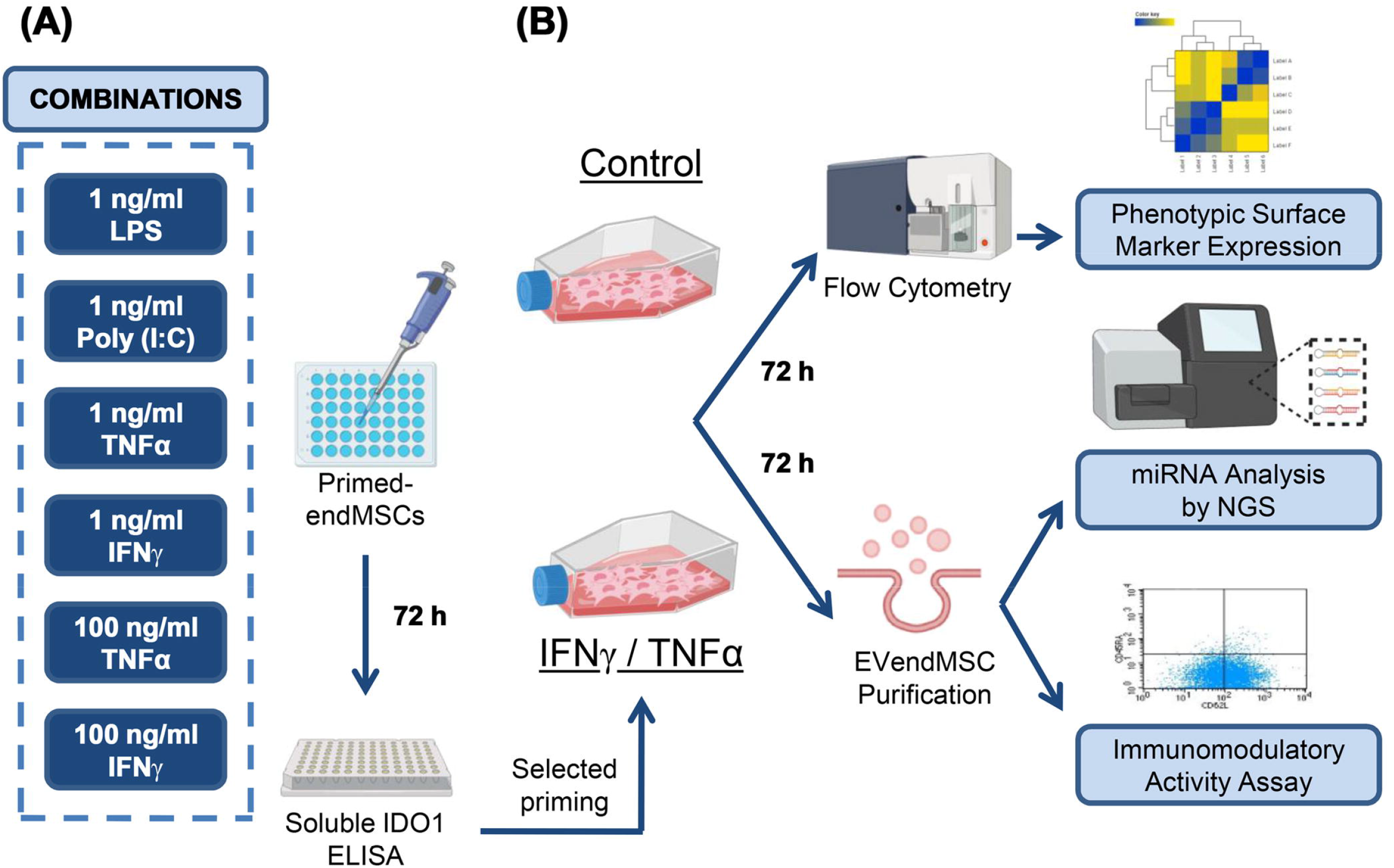
Graphic overview of key experimental procedures. (A) IFNγ, TNFα, LPS, and Poly (I:C) were used at different concentrations, single or in combination, to prime *in vitro* cultured endMSCs. The optimal priming strategy was identified by ELISA in terms of IDO1 release. (B) In accordance with ELISA results, endMSCs were primed with 100 ng/ml IFNγ/TNFα, and the phenotypic profile of primed and control endMSCs was determined by flow cytometry. EndMSC and endMSC* collected medium were concentrate to get secretomes. The molecular profile of secretomes was analyzed by Next Generation Sequencing (NGS). In addition, the immunomodulatory activity of control S-endMSCs or S-endMSCs* was analyzed in a lymphocyte activation assay. Images have been created with BioRender (https://app.biorender.com/).

### Priming of endMSCs and IDO1 quantification by ELISA

Different lineages of *in vitro* cultured endMSCs (n = 3) were primed using a combination of recombinant pro-inflammatory cytokines and Toll-like receptors (TLR) ligands: IFNγ (Miltenyi Biotec Inc, San Diego, CA, USA) at 1 ng/ml and 100 ng/ml, TNFα (Miltenyi Biotec Inc) at 1 ng/ml and 100 ng/ml, LPS (Sigma, St. Louis, MO, USA) at 1 ng/ml and Poly (I:C) (Miltenyi Biotec Inc) at 1 ng/ml. At day 3, supernatants from *in vitro* primed endMSCs were collected to evaluate the release of IDO1. According to the manufacturer’s instructions, the IDO1 levels were quantified by ELISA (R&D SYSTEMS, Minneapolis, USA).

### Phenotypic surface markers and comparison of endMSCs and endMSCs*

The phenotypic analysis of the cells was performed by flow cytometry using a panel of human monoclonal antibodies (Supplementary table 1). The endMSCs at passages 5–7 and 80% confluence, were *in vitro* cultured devoid of recombinant IFNγ and TNFα (endMSCs, n = 3) or with 100 ng/ml IFNγ and 100 ng/ml TNFα for 72 h (endMSCs*, n = 3). The endMSCs and endMSCs* were detached from tissue culture flasks using PBS containing 0.25% trypsin and incubated with different antibodies for 30 min at 4 °C in PBS with 2% of FBS. Cells were then washed, resuspended in PBS and acquired in a FACSCalibur™ cytometer (BD Biosciences, San Jose, CA, USA) equipped with CellQuest software (BD Biosciences). Detached cells were incubated with the Fluorescence Minus One Control (FMO control) to properly compensate the flow cytometry data. Mean fluorescence intensities (MFI) and standard deviations (SD) of positive and negative populations were taken into consideration to calculate the Stain Index (SI) as follows: MFI (positive population) – MFI (negative population) / 2 × SD (negative population).

Phenotypic profiles of endMSCs and endMSCs* were compared. Significant differentially expressed proteins underwent enrichment and biological pathway analyses with Cytoscape (version 3.7.2) (52), which includes the Gene Ontology Resource1 and the Reactome Pathway Database2.

### Isolation, purification, and quantification of Secretomes

The endMSCs (n = 3) and endMSCs* (n = 3) (obtained as mentioned above) at passages 6–12 and 80% confluence were used for secretome collection. Cell culture medium (with or without IFNγ and TNFα) was replaced by DMEM, 1% penicillin/streptomycin, 1% glutamine and 1% insulin-transferrin-selenium (Thermo Fisher Scientific, MA, USA), after rinsing with PBS. Cells were then cultured during 72 h at 37 °C and 5% CO_2,_ conditioned media were collected for the concentration of secretomes. Cells were detached with trypsin and counted with Neubauer chamber. Viability was 168 evaluated though Trypan blue staining (Thermo Scientific).

Condition media from endMSCs and endMSCs* were centrifuged first at 1000 × g 10 min 4 °C, then at 5000 × g 20 min 4 °C. Pellets were discarded and the supernatants were filtered through 0.45 µm and 0.22 µm mesh (Fisher Scientific, Leicestershire, UK) to eliminate dead cells and debris. To concentrate the condition media, up to 15 ml filtered supernatants were ultra-filtered through a 3 kDa MWCO Amicon® Ultra device (Merck-Millipore, MA, USA) by centrifugation at 4000 x g for 1 h at 4 °C. The concentration of proteins from enriched secretomes was quantified by Bradford assay (Bio Rad Laboratories, Hercules, CA). Finally, the secretomes from endMSCs (S-endMSCs) and endMSCs* (S-endMSCs*) were stored at -20°C for further analyses.

### miRNAs analysis by Next Generation Sequencing

miRNA sequencing experiments were performed at QIAGEN Genomic Services (Hilde, Germany). Total RNA was isolated from 1 ml of each concentrated sample, using the exoRNeasy Serum/Plasma Kit (QIAGEN) according to manufacturer’s instructions. During the sample preparation, QC Spike-ins were added as quality control. A total of 5 μl of purified RNA tagged with adapters containing a Unique Molecular Index (UMIs) was converted into cDNA NGS libraries using the QIAseqmiRNA Library Kit (QIAGEN). cDNA was amplified in 22 cycles of PCR and purified. Library preparation QC was performed using Bioanalyzer 2100 (Agilent). Libraries were pooled in equimolar ratios, based on quality of the inserts and the concentration measurements. They were then quantified by quantitative polymerase chain reaction (qPCR) and sequenced on a NextSeq500/550 System (Illumina, San Diego, CA, USA) according to the manufacturer instructions. Raw data was converted to FASTQ files for each sample using the bcl2fastq software (Illumina, Inc.). Quality control of raw sequencing data was checked using the FastQCtool^3^. The data analyses, which included filtering, trimming, mapping, quantification, and normalization, were carried out using CLC Genomics Workbench (version 20.0.2) and CLC Genomics Server (version 20.0.2). The human genome version GRCh38 was used as a reference database to annotate the miRNAs. The read sets were aligned to the reference sequences from miRbase (version 22). The differential expression between S-endMSCs and S-endMSCs* was evaluated through the EDGE (Empirical analysis of Differential Gene Expression) analysis within the CLC bio.

Using Benjamini-Hochberg FDR corrected *p* values, differentially expressed miRNAs at significance level of .05 (FDR) were selected for unsupervised analysis (principal component analysis -PCA-, clustering, and heat map) using ClustVis (version 2.0)^4^. Enrichment analysis of these miRNAs was performed using miRNet (version 2.0)^5^ and TAM (version 2.0)^6^. Moreover, significantly overexpressed miRNAs in S-endMSCs* were submitted to a miRTargetLink^7^ analysis to determine the human target genes of these miRNAs. Results were filtered to strong experimental evidence. Only genes included into the Gene Ontology category of *Inflammation Response* (GO:0006954) were taken into consideration. Enrichment analysis of these genes was performed using FunRich: Functional Enrichment analysis tool (version 3.1.3) (53,54). Target genes were sub-classified in four categories: *Innate Immune Response* (GO:0045087), *Adaptive Immune Response* (GO:0002250), *Positive Regulation of Inflammatory Response* (GO:0050728), and *Negative Regulation of Inflammatory Response* (GO:0050729). The datasets discussed in this publication have been deposited in Sequence Read Archive (SRA) data with the accession number PRJNA664968^8^.

### Immunomodulatory assays on *in vitro* stimulated peripheral blood lymphocytes

Human peripheral blood lymphocytes (PBLs) from a healthy donor were stimulated using the T Cell Activation/ Expansion Kit (MACS,Miltenyi Biotec, USA) according to manufacturer instructions. Briefly, PBLs in RPMI-1640 medium supplemented with T Cell Activation/ Expansion Kit and 10% FBS were cultured in U-bottom 96 wells plate at 4 × 10^5^ cells per well. S-endMSCs and S-endMSCs* were added to PBLs at 20, 40, and 80 µg/ml, in accordance with protein concentrations resulting from the Bradford assay. At day 3 and day 6, *in vitro* stimulated PBLs were analyzed by flow cytometry. PBLs without stimulation were used as negative control. *In vitro* stimulated PBLs without secretomes were used as positive controls. For flow cytometry analyses, PBLs were collected and incubated for 30 min at 4 °C with fluorescence-labeled human monoclonal antibodies against: CD4, CD8, CD45RA, and CD62L (BD Biosciences), in the presence of PBS containing 2% of FBS. Cells were then washed and resuspended in PBS. The flow cytometric analysis was performed on CD4+ T cells, and CD8+ T cells, and the percentage of gated CD45RA-/CD62L-cells allowed us to quantify the percentage of effector-memory cells.

### Statistical analysis

Data were statistically analyzed with GraphPad Prim (version 8.0) using paired *t* test for variables with parametric distribution and Wilcoxon test for non-parametric data. The *p* ≤ .05 were considered statistically significant.

## Results

### Characterization of *in vitro* isolated and expanded endMSCs

A more detailed description of endMSC characterization can be found in our previous studies (38,39,51). Briefly, our endMSCs were plastic-adherent when maintained in standard culture conditions and their differentiation toward the adipogenic, chondrogenic, and osteogenic lineages was demonstrated by specific staining. The phenotypic analysis of *in vitro* expanded endMSCs revealed a positive expression for the surface markers CD44/CD73/CD90/CD105, and negative for CD14/CD20/CD34/CD45/CD80/HLA-DR. The cell lines used for this study complied with the Minimal criteria for defining multipotent mesenchymal stromal cells (11).

### Effect of endMSCs priming on IDO1 release

In order to enhance the immunomodulatory capacity of secretomes from endMSCs, different pro-inflammatory cytokines (TNFα, IFNγ) and TLR ligands (LPS, Poly (I:C)) were evaluated under *in vitro* conditions. The release of IDO1 by endMSCs was used as a primary biomarker for the immunomodulatory potential (55). Results obtained by ELISA firstly demonstrated an increase in IDO1 release when endMSCs were primed with 100 ng/ml of IFNγ. The significant increase was also observed when 100 ng/ml of IFNγ were combined with LPS and/or Poly (I:C) at 1 ng/ml. In the case of IFNγ at low concentrations (1 ng/ml), alone or in combination, the release of IDO1 was not increased when compared to control conditions.

Interestingly, the combination of 100 ng/ml IFNγ and 100 ng/ml TNFα achieved the highest level of IDO1 release (Figure 2). These results demonstrated a synergistic effect of IFNγ and TNFα on the immunomodulatory potential of endMSCs. Hence, endMSCs under these *in vitro* culture conditions were characterized in terms of surface marker expression, and their secretomes were concentrated and characterized.

**Figure 2.**
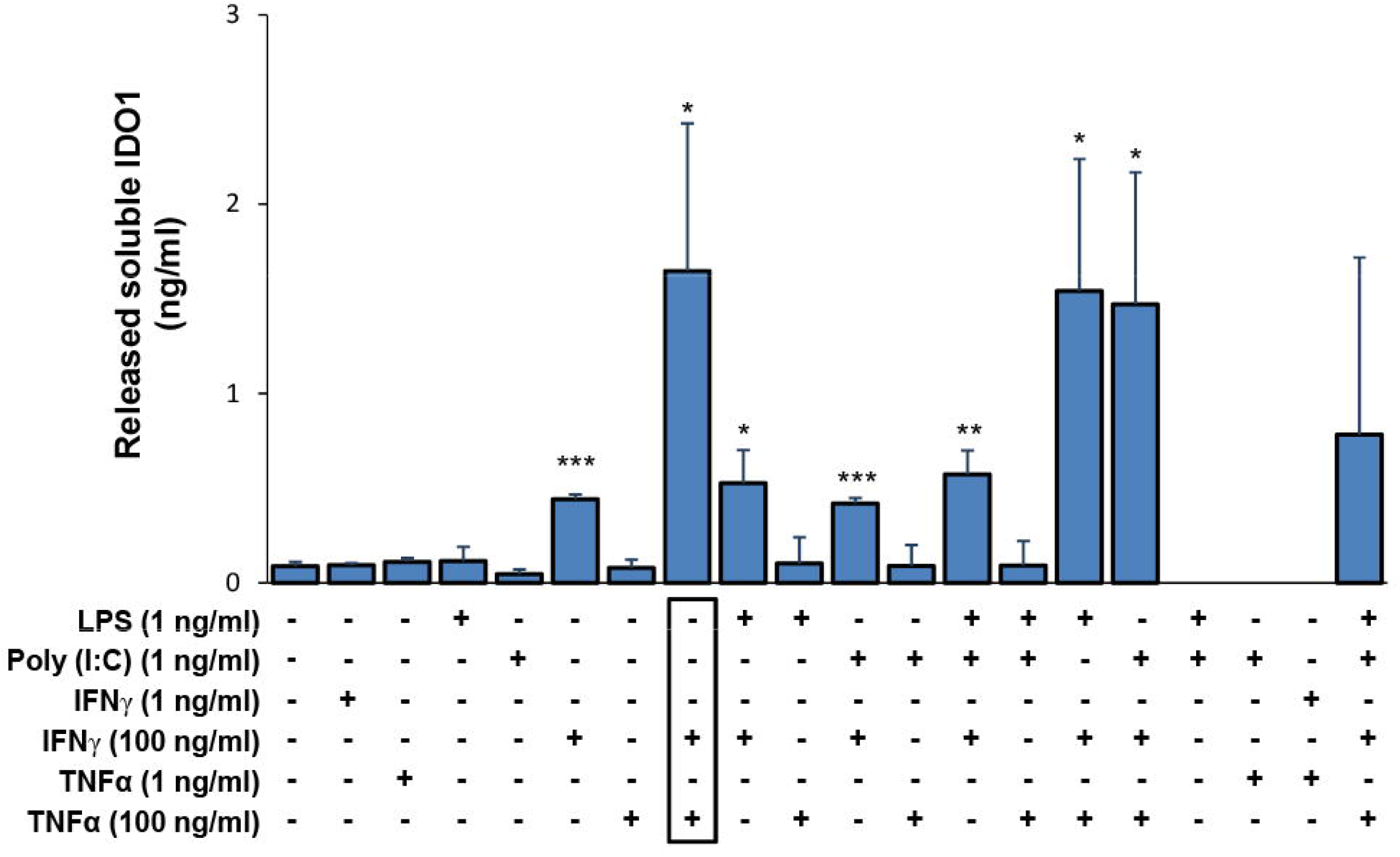
Release of soluble IDO1 by primed endMSCs. endMSCs (n = 3) were primed with several combinations and concentrations of proinflammatory cytokines: IFNγ and TNFα, and TLR ligands: LPS and Poly (I:C). Levels of released IDO1 by the cells were tested by ELISA. A paired *t*-test was used to compare IDO1 levels in control endMSCs with the different priming conditions. Error bars represent the standard deviations of data. Asterisks indicate statistically significant differences: \**p* < .05, \*\**p* < .01, *** *p* < .001.

### Phenotypic profile of surface markers on endMSCs

endMSCs (n = 3) and endMSCs* (n = 3) phenotype was analyzed by flow cytometry using a large panel (n = 40) of monoclonal antibodies toward various cell surface markers (Supplementary Table 1). Our phenotypic analysis revealed a statistically significant difference in 9 out of 40 surface markers between endMSCs and endMSCs* (Figure 3A).

**Figure 3.**
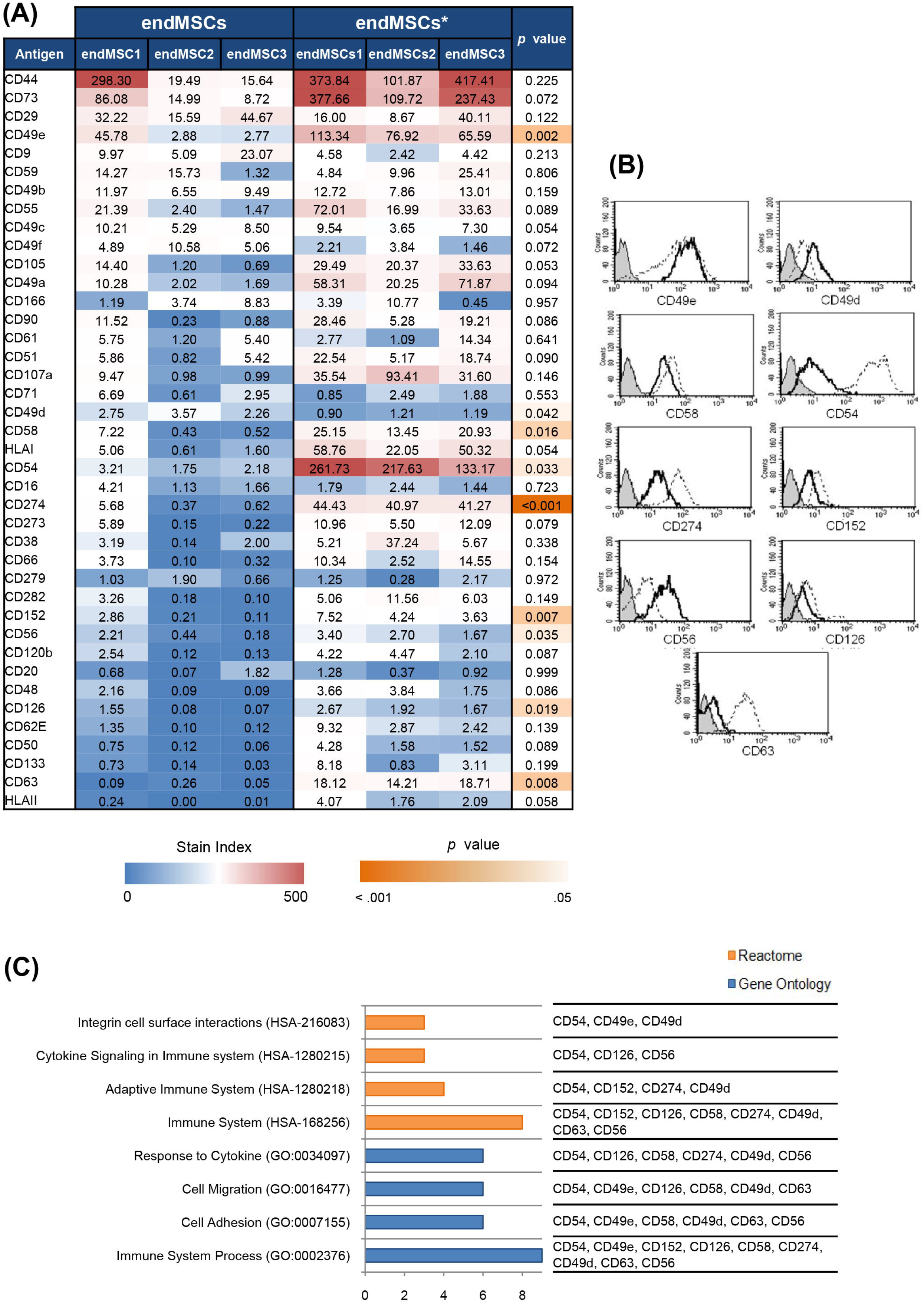
Phenotypic surface marker expression on endMSCs. Analysis of surface markers in control endMSCs (n = 3) and endMSCs* (n = 3) using 100 ng/ml IFNγ and 100 ng/ml TNFα at 3 days. (A) The expression level of 40 cell surface markers (Stain Index (SI) value) were illustrated in a heat map. To compare control endMSCs and endMSCs*, a paired *t*-test was carried out and *p* < .05 were considered statistically significant. The color scale for the SI values gives the highest expression values (red) and the lowest (blue); the orange color scale indicates the grade of signification, being the most significant ones in dark orange. (B) Representative histograms of statistically different markers are shown. The expression in endMSCs is represented by a black lined histogram and in endMSCs* by a discontinuous line. The opaque gray histogram corresponds to the fluorescence negative control. (C) Reactome and Gene Ontology enrichment analyses of statistically significant different markers were performed by adjusting *p* value by Benjamini-Hochberg FDR correction < .01. Graph bars represent the number of surface markers included into each category of Reactome (orange) and Gene Ontology (blue).

The following markers were significantly increased in endMSCs*: CD49e, CD54, CD56, CD58, CD63, CD126, CD152, and CD274. Uniquely, CD49d was significantly reduced in IFNγ/TNFα-primed cells (Figure 3B). Finally, differentially expressed antigens were classified according to Gene Ontology and Reactome Pathways. Our results revealed the classification of these markers in *Immune System Process* (GO:0002376) (CD54, CD49e, CD152, CD126, CD58, CD274, CD49d, CD63, CD56), *Cell Adhesion* (GO:007155) (CD54, CD49e, CD58, CD49d, CD63, CD56), *Cell Migration* (GO:0016477) (CD54, CD49e, CD126, CD58, CD49d, CD63), and *Response to Cytokine* (GO:0034097) (CD54, CD126, CD58, CD274, CD49d, CD56). The analysis by Reactome Biological Pathways demonstrated a classification in *Immune System* (HSA-168256) (CD54, CD152, CD126, CD58, CD274, CD49d, CD63, CD56), *Adaptive Immune System* (HSA-1280218) (CD54, CD152, CD274, CD49d), *Cytokine Signaling in Immune system* (HSA-1280215) (CD54, CD126, CD56), and *Integrin cell surface interactions* (HSA-216083) (CD54, CD49e, CD49d) (Figure 3C).

### Effects of IFN_γ_/TNF_α_ priming in S-endMSCs microRNAome

Particle size of secretomes was determined by nanoparticle tracking analysis. Particle diameter was 153.5 ± 63.05 nm, as already reported in one of our previous studies (39). A microRNAome NGS analysis was performed on concentrated S-endMSCs (n = 3) and S-endMSCs* (n = 3). A total of 628 miRNAs were identified from S-endMSCs. Supplementary Table 2 shows the expression in counts per millions (CPM) of top ten abundant miRNAs in the secretomes. Additionally, the comparison of the normalized and filtered expression values of CPM revealed that 40 of them (6.37%) were significantly different when S-endMSCs and S-endMSCs* were compared (*p* value adjusted by Benjamini-Hochberg FDR correction ≤ .05). The top ten of miRNAs with the highest fold change were hsa-miR-155-5p, hsa-miR-361-3p, hsa-miR-376a-3p, hsa-miR-424-3p, hsa-miR-27a-3p, hsa-miR-210-3p, hsa-miR-21-3p, hsa-miR-490-3p, hsa-miR-26a-2-3p, and hsa-miR-181a-5p (Supplementary Table 3).

Our results demonstrated that, 25 out of 40 significantly differentially expressed miRNAs (62.5%) were up-regulated in S-endMSCs*. These 25 miRNAs were selected for further analyses (unsupervised analysis and Gene Ontology enrichment analysis). To address the differences in miRNAome profile between the S-endMSCs* and S-endMSCs, unsupervised principal component analysis was used to study the potential clusters of these S-endMSCs based on the detected miRNAs. The result of the PCA indicated that each condition exhibited a unique miRNA profile, allowing a separation into well differentiated clusters (Figure 4A). The heat map-based unsupervised hierarchical clustering analysis showed the differences in these 25 miRNAs (Figure 4B) and corroborated the PCA analysis.

**Figure 4.**
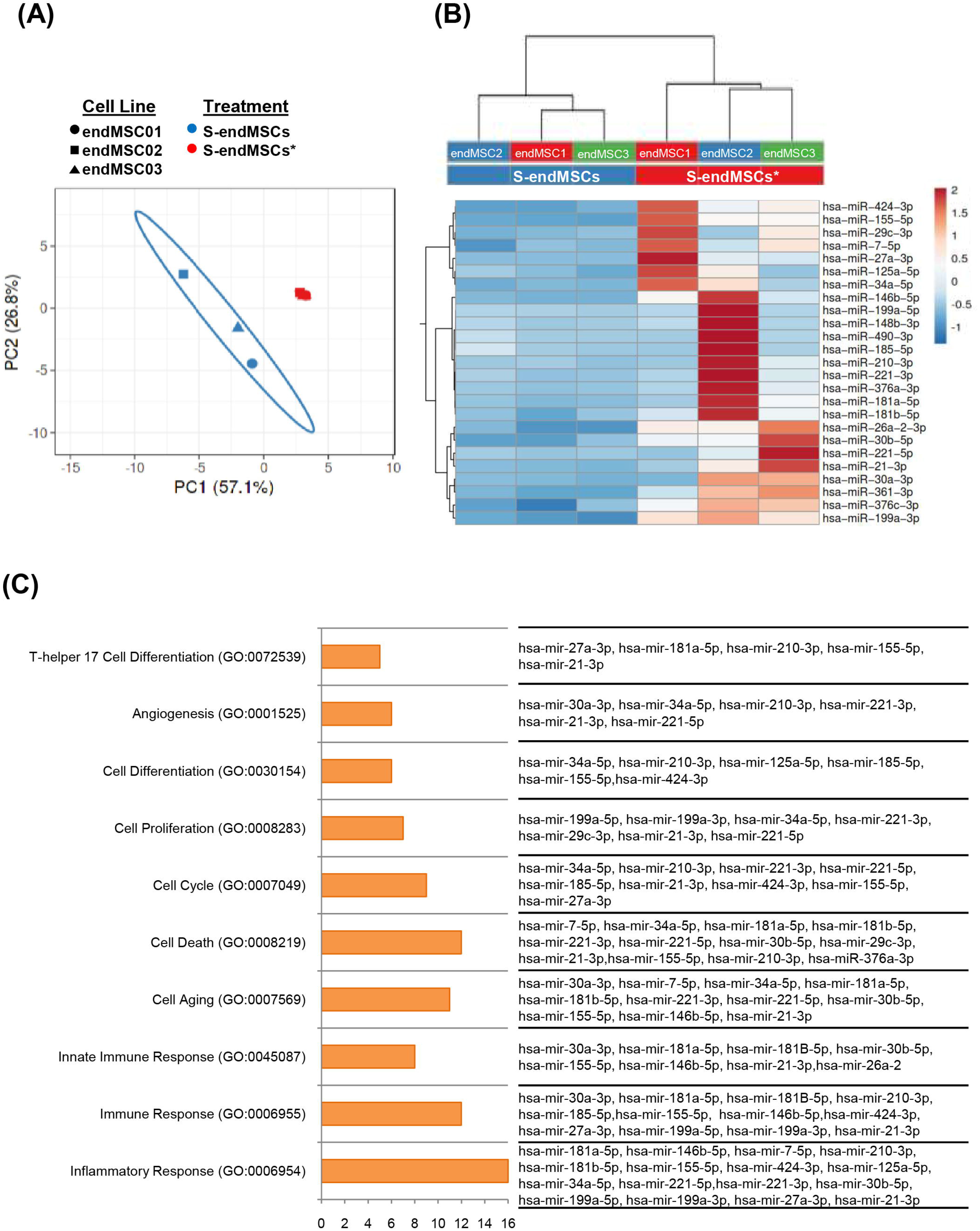
Effects of IFN_γ_/TNF_α_ priming on S-endMSCs microRNAome. A selection of the 25 miRNAs which showed a significant increase in S-endMSC*, were used for further analyses. (A) Principal Component Analysis (PCA) plots. Score plot for PC1 (57.1% variance explained) vs. PC2 (26.8% variance explained). (B) Hierarchical Clustering of secretomes (S-endMSC1, S-endMSC2, S-endMSC3) and the different conditions (basal red and primed in blue) together with the heat map corresponding to the selected miRNA expressions. (C) Enrichment analysis revealed a significant implication of the selected miRNAs in Gene Ontology categories *(p* value adjusted by Benjamini-Hochberg FDR correction < .01). Graph bars represent the number of miRNAs in each category.

Gene Ontology enrichment analysis was performed to classify these up-regulated miRNAs. For this analysis, miRNet and TAM 2.0 software were used. Several biological processes turned out to be enriched, being the most relevant the terms: *Inflammatory Response* (GO:0006954) (64%), *Immune Response* (GO:0006955) (48%), *Cell Death* (GO:0008219) (48%), *Cell Aging* (GO:0007569) (44%), *Cell Cycle* (GO:0007049) (36%), *Innate Immune Response* (GO:0045087) (32%), *Cell Proliferation* (GO:0008283) (28%), *Cell Differentiation* (GO:0030154) (24%), *Angiogenesis* (GO:0001525) (24%), and *T-helper 17 Cell Differentiation* (GO:0072539) (20%).

### miRNA Target Prediction in Inflammatory Response

According to microRNAome results, the up-regulated miRNAs were subsequently analyzed for determining the miRNA target genes using human miRTargetLink. Only miRNA-target interactions with strong experimental evidence were included. Our results demonstrated that, 105 genes were identified as miRNA targets (a total of 6 miRNAs were excluded because of a lack of connections). An enrichment analysis, carried out with FunRich, revealed that 22 genes out of 105 were included into Gene Ontology category of *Inflammation Response* (GO:0006954). These genes were sub-classified in four categories: *Innate Immune Response* (GO:0045087), *Adaptive Immune Response* (GO:0002250), *Positive Regulation of Inflammatory Response* (GO:0050728), and *Negative Regulation of Inflammatory Response* (GO:0050729). Some genes belonged to more than one category. The classification showed that 8 target genes were categorized in *Innate Immune Response* (*CD44, IFNG, PIK3CG, CSF1R, ICAM1, XIAP, APOE, NFKB1*), 6 target genes were categorized in *Positive Regulation of Inflammatory Response* (*PTGS2, IFNG, PIK3CG, ETS1, CEBPB, EGFR*), 5 target genes were categorized in *Negative Regulation of Inflammatory Response* (*IGF1, ETS1, APOE, KLF4, NFKB1*), and 4 target genes were categorized in *Adaptive Immune Response* (*BCL6, IFNG, PIK3CG, ICAM1*). Moreover, the genes: *ATM, KIT, FOS, SELE, SMAD1, NOTCH1*, and *HIF1A* were not included in these sub-categories. Finally, a Protein-miRNA Interaction Networks with 35 nodes and 51 edges were built using Cytoscape software (Figure 5).

**Figure 5.**
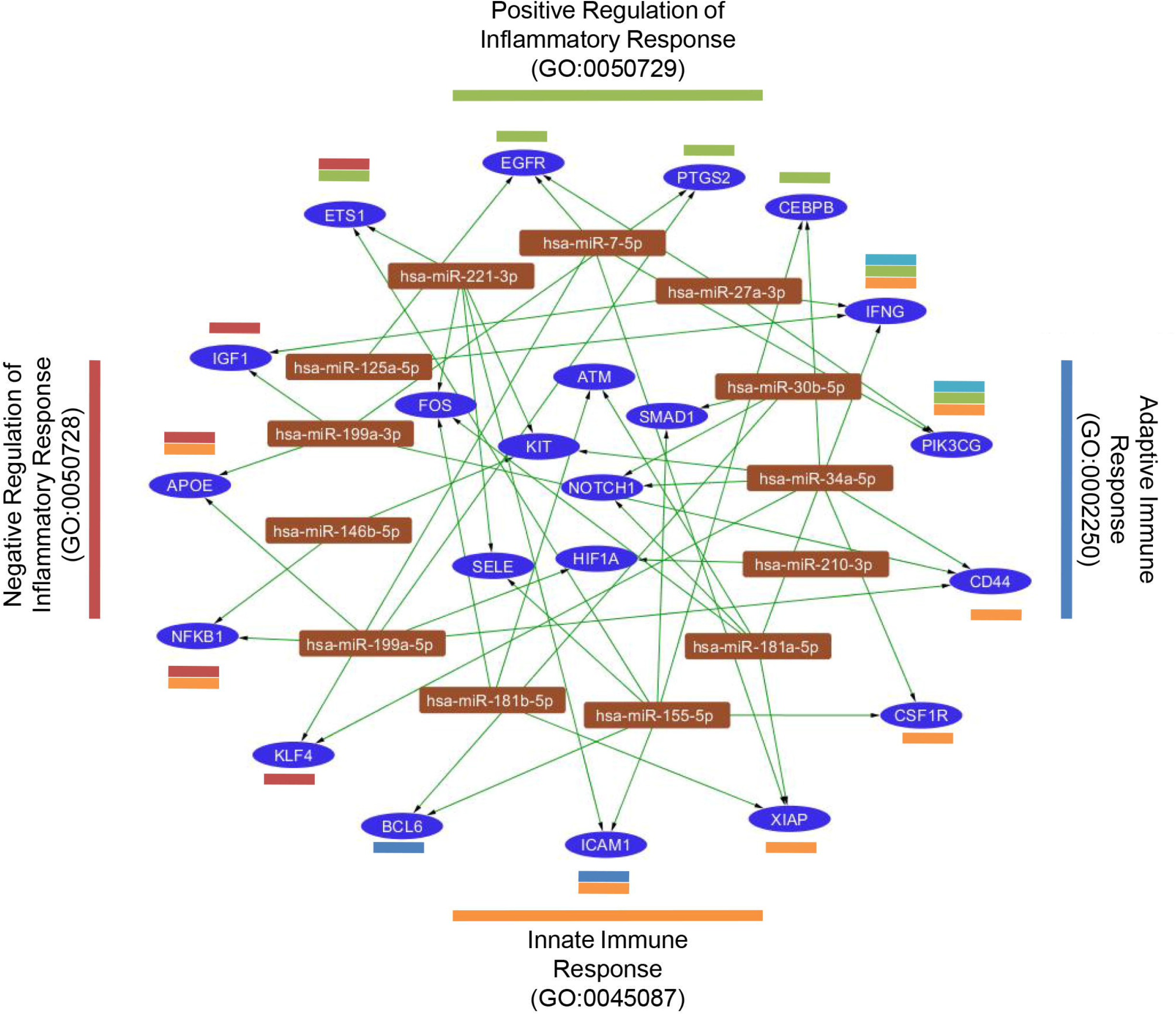
Predicted target genes for miRNAs. Target genes of significantly increased miRNAs from S-endMSC*. Target genes classified into the category *Inflammatory Response category* (GO:0006954) were sub-classified into four sub-categories (*Innate Immune Response* (orange), *Adaptive Immune Response* (blue), *Positive Regulation of Inflammatory Response* (green), and *Negative Regulation of Inflammatory Response* (red). Interaction network of gene targets (dark blue ellipses) and miRNAs (brown rectangles) were illustrated using Cytoscape Software.

### Immunomodulatory assay of secretomes against *in vitro* stimulated lymphocytes

In order to evaluate the immunomodulatory effect of extracellular vesicles against *in vitro* stimulated lymphocytes, PBLs from a healthy donor were stimulated with anti-CD2, anti-CD3, and anti-CD28 beads. This stimulation partially mimics antigen-mediated activation and trigger the differentiation of CD4+ and CD8+ T cells towards effector-memory T cells (CD45RA-/CD62L-).

Lymphocyte activation assays were performed co-culturing PBLs in the presence of secretomes at 20, 40, and 80 µg/ml for 3 and 6 days. The most relevant results were obtained at day 6 with a secretome concentration of 40 µg/ml. The phenotype of *in vitro* stimulated T cells after co-culture with secretome 40 µg/ml at 6 days is shown in Figure 6. As expected, S-endMSCs* triggered a significant decrease in the percentage of CD4+ effector memory T-cells when compared to *in vitro* stimulated PBLs devoid of secretomes (*p* = .0428) and to S-endMSCs (*p* = .0313) (Figure 6A and 6C). Additionally, S-endMSCs* produced a significant decrease in the percentage of CD8^+^ effector memory T-cells when compared to *in vitro* stimulated PBLs (*p* = .0043) and to S-endMSCs (*p* = .0283) (Figure 6B and 6D). The percentage of effector-memory T cells was determined as the percentage of CD45RA-/CD62L-cells on FSC/SSC-gated cells.

**Figure 6.**
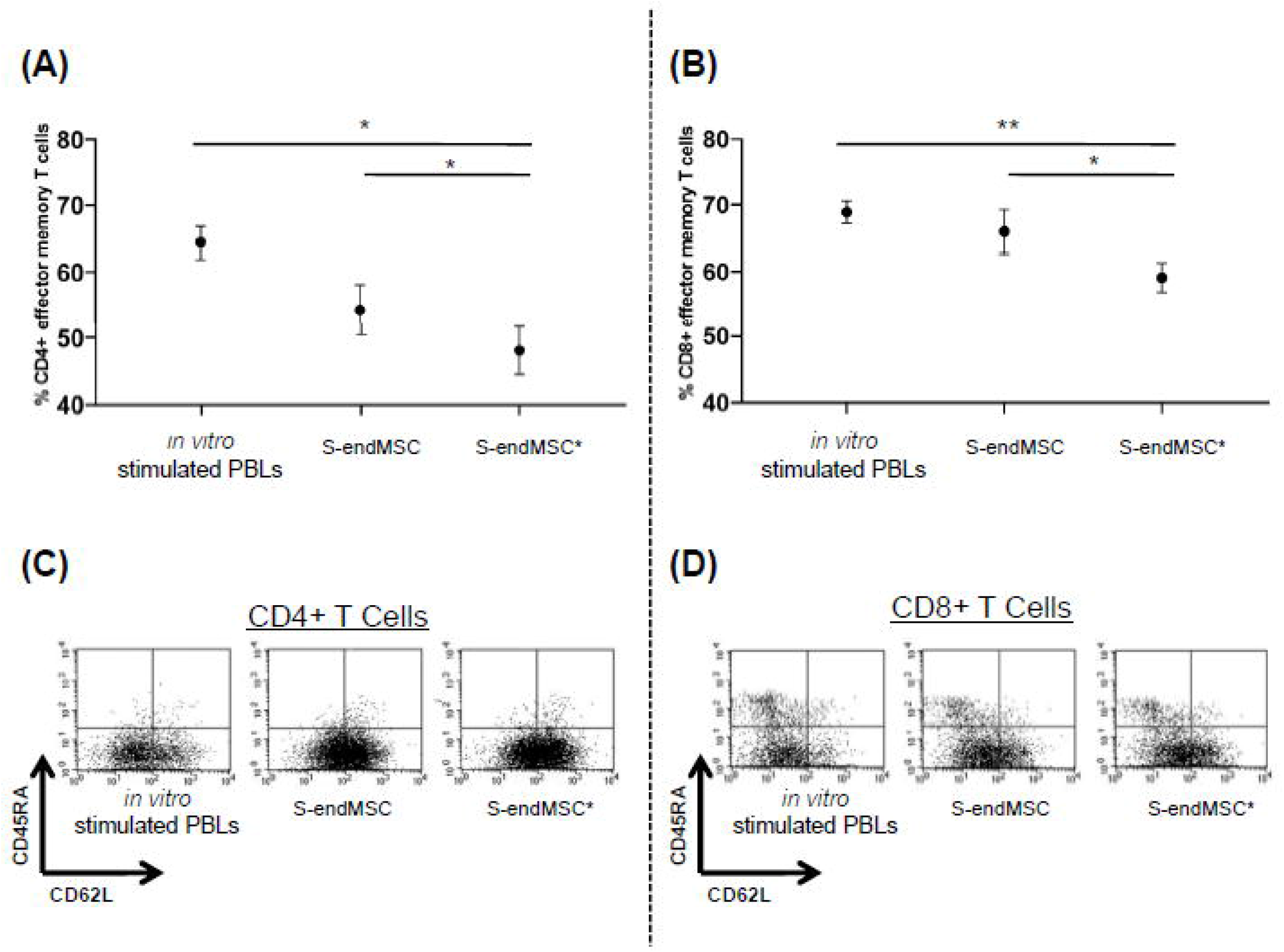
Lymphocyte activation assays for the evaluation of Secretomes. PBLs from a healthy donor were stimulated and simultaneously co-cultured with S-endMSCs (n = 3) and S-endMSCs* (n = 3) at 40 µg/ml for 6 days. (A) Percentage of CD4+ and (B) percentage of CD8+ effector memory T cells (CD45RA-/CD62L-). Data from non-stimulated PBLs is not shown. Values represent the mean ± SD of independently performed experiments. Data were statistically analyzed using a paired *t*-test and *p* value was considered significant at < .05. *Statistically significant differences (*p* < .05). **Statistically significant comparison (*p* < .005). A representative dot plot of each condition is illustrated: (C) CD4+ and (D) CD8+ T cells. The gates for CD45RA and CD62L were drawn according to negative controls.

## Discussion

Secretome derived from MSCs have become a promising tool for the regulation of adverse inflammatory events and to support regenerative processes. Among secretome sources, endometrial MSCs derived from menstrual blood (endMSCs) present many advantages: they can be obtained by non-invasive procedures, from multiple donors and without ethical concerns. Isolating and expanding these cells is easy and feasible, guaranteeing high growth rates in relatively short time (56). In earlier studies, we have demonstrated that their secreted vesicles had an immunomodulatory effect on CD4+ T cells (38). In addition, proteomic/genomic analyses revealed the presence of proteins and miRNAs involved in immunomodulatory pathways (39).

Here we aimed to improve *in vitro* culture conditions to enhance the immunomodulatory capacity of secretomes from endMSCs, and hence, their therapeutic effectiveness. According to bibliography, different priming -or licensing-strategies to enhance the immunomodulatory capacity of MSCs have been proposed (57). Pro-inflammatory cytokines, hypoxia, biological or chemical agents have been successfully used to enhance the immunomodulatory capacity against immune cells such as T cells, B cells, NK cells, neutrophils or dendritic cells (18,19). It should be pointed out that most of these results were obtained under *in vitro* conditions and several concerns about using these licensed cells in clinical settings have arisen. As an example, they may allow tumor development of pre-existing malignant cells (58).

Based on this idea, our first sets of experiments were focused on the optimization of priming conditions. IFNγ, TNFα, Poly (I:C), and LPS (separately or in combination) were used. For these experiments, IDO1 secretion was considered an “immunomodulatory biomarker” and allowed us to compare the efficiency of different priming conditions. It is important to note that, this enzyme has been also considered by other authors as a “potency marker” for MSC-mediated immune suppression (59). Our experimental conditions revealed that IFNγ itself enhanced the release of IDO1, which is in agreement with *in vitro* studies using adipose-derived MSCs (60). Additionally, it has been recently demonstrated that IFNγ causes chromatin remodeling at the IDO1 promoter increasing the mRNA levels (41). Our results were remarkable when IFNγ was combined with TNFα. These cytokines showed a synergistic effect and evidently increased the release of IDO1 from endMSCs. Although the combination of these two cytokines is not new, and it has been already used to induce the immunomodulatory activity of MSCs (49,50), this the first report using MSCs from menstrual blood.

Using IDO1 as immunomodulatory biomarker, endMSC priming conditions were optimized. Hence, the next set of experiments was conducted to characterize the phenotype of IFNγ/TNFα-primed cells. In this analysis, a large panel of surface markers (n = 40) was analyzed to compare endMSCs* and endMSCs. As expected, endMSCs were positive for CD44, CD73, CD90, and CD105 with low levels of MHC class I molecule and negative expression of MHC class II. The statistics and paired comparisons revealed that 9 out of 40 surface markers were significantly increased in endMSCs*. Most of these markers were classified by Gene Ontology and Reactome in the categories *Immune System Process* (GO:0002376) and *Immune system* (R-HSA-168256) respectively.

It is important to discuss the significant differences observed in some of these molecules. NCAM-1 (CD56) is an adhesion molecule expressed in bone marrow-derived MSCs and associated with their migration and homing capacity (61). The surface molecule ICAM-1 (CD54) in MSCs promotes MSC homing to the focus of inflammation and immune organs (62) and its increased expression has been correlated with an immunomodulatory effect on dendritic cells (63). The expression of LFA3 (CD58) has been also described in bone marrow-derived MSCs and interacts with their ligands on T cells (64). Considering that preclinical and clinical studies have demonstrated that the homing/migration of systemically administered MSCs is very low (66), the increased expression of adhesion molecules ICAM-1, CD58, and NCAM-1 in IFNγ/TNFα-primed cells may improve homing and migration capacity to a target tissue.

Apart from adhesion molecules, our phenotypic analysis was also focused on immune checkpoint receptors such as CTLA4 (CD152) and PD-L1 (CD274). Here we demonstrate a significant increase of the inhibitory molecule CTLA4 (CD152) under IFNγ/TNFα priming. The expression of this protein has been previously described in the membrane of bone marrow-derived MSCs and its soluble form is released under hypoxic conditions (67). In our study, we also found a significant increase of CTLA4 in endMSCs*, which may have an immunomodulatory role by blocking CD28 co-stimulation on T cells. Similarly to CTLA4, we found that PD-L1 (CD274) was also increased after IFNγ/TNFα priming. This result is coincident with previous studies where bone marrow-derived MSCs express and secrete PD-L1 (CD274), that can be inducible under IFNγ and TNFα (68). The surface expression and inflammatory-mediated induction of CTLA4 and PD-L1 suggests that these molecules could be involved in the modulation of T cells and peripheral tolerance.

Finally, IL6R (CD126) emerged from our phenotypic analysis. endMSCs as adipose-derived MSCs (69) did not express -or had a very low expression of-CD126. Previous studies have demonstrated that CD126 cannot be induced in adipose-derived MSCs (using IFNγ or TGFβ) (69). In contrast, in our experimental conditions we found a significant increase of CD126. The expression of IL6/IL6R has been correlated with the osteogenic (70) and adipogenic (71) differentiation of bone marrow-derived MSCs. Here we assume that the increase of CD126 may also reflect an increased susceptibility of endMSCs to IL6-mediated inflammation.

The second aim of this study was to analyze the secretomes from endMSCs*. As previously discussed, our group recently demonstrated the involvement of miRNAs in immunomodulatory pathways (39). Therefore, we hypothesized that IFNγ/TNFα may trigger the release of miRNAs that target genes from inflammation and innate/adaptive immune responses. The analysis by NGS allowed us to identify a set of significantly increased (n = 25) and significantly decreased (n = 16) miRNAs following IFNγ/TNFα-priming. According to our previous studies, three of them were also significantly expressed when using IFNγ alone for priming endMSCs: hsa-miR-146b-5p, hsa-miR-376c-3p, and hsa-miR-490-3p (39). Similarly, here we confirm our previous findings which demonstrated that hsa-miR-143-3p, hsa-miR-199a-3p, hsa-let-7b-5p, hsa-miR-21-5p, hsa-miR-16-5p, hsa-let-7a-5p, and hsa-let-7f-5p, are the most abundant miRNAs in secretomes (39).

It is interesting to note that the PCA of NGS results evidenced the significant differences among secretomes under standard culture conditions and secretomes from endMSCs*. These results highlight the importance of cell culture conditions in the secretome release, and how inflammatory priming conditions determine the miRNAs content. Although some of these miRNAs may deserve a long discussion, we have focused our interest on those miRNAs which are significantly increased and target inflammatory-related genes.

The miRTargetLink analysis (72,73) allowed us to identify multiple query nodes on significantly increased miRNAs. The increase of hsa-miR-155-5p (log fold change = 5.05 and *p* = .00009) is especially relevant, since it targets genes involved in *Innate/Adaptive Immune Response*, such as *CSF1R, ICAM1* or *BCL6*. This result agrees with a recent study from E. Ragni et al., who demonstrated that miR-155-5p is overexpressed in extracellular vesicles from IFN-γ-primed adipose-derived MSCs (74). Similarly, an enhancement of miR-155-5p was previously reported in bone marrow-derived MSCs under IFNγ/TNFα stimulation, with the authors suggesting that this miRNA could inhibit the immunosuppressive capacity of MSCs by reducing the iNOS production (75). In contrast, a recent *in vitro* study using inhibitors of miR-155-5p in bone marrow-derived MSCs from rats demonstrated that miR-155-5p enhanced the differentiation of T cells towards Th2 and Treg cells (76). Based on our *in vitro* assays using *in vitro* stimulated T cells co-cultured with secretomes, we believe that miR-155-5p could be involved in the inhibition of T cell activation. Obviously, future studies must be conducted to validate this hypothesis.

The increase of hsa-miR-27a-3p was also significant in S-endMSCs* (log fold change = 4.35 and *p* = .000005). This miRNA has been recently described in extracellular vesicles derived from MSCs, and more importantly, it has been found to be transferred from vesicles to macrophages promoting M2 macrophage polarization (77). Furthermore, according to miRTargetLink, the gene *IFNG* is a target of hsa-miR-27a-3p, and this miRNA has been found to be involved in the regulation of *IRAK4*, a promoter of *NF-*κ*B* (78), which regulate inflammatory and immune genes.

In the case of hsa-miR-21-3p and hsa-miR-490-3p, these miRNAs were abundantly expressed and significantly increased in S-endMSCs under IFNγ priming (39). Similarly, hsa-miR-424-3p has been found to be increased in IFNγ-primed umbilical cord-derived MSCs (79). Regarding hsa-miR-185-5p, a recent study using EVs from MSCs have demonstrated that the enrichment of these vesicles with this miRNA alleviate the inflammatory response and reduce cell proliferation (80).

Altogether, data analysis by NGS revealed a myriad of miRNAs which are increased and decreased in S-endMSCs*. According to Gene Ontology, the targets for these miRNAs were found to be involved in the *regulation of inflammatory response*, so it is expected that the transference of these miRNAs to target cells may have an impact in the behavior of immune cells, and hence, in the TH1/TH2 balance. Our functional studies using *in vitro* stimulated lymphocytes (that partially mimic an antigen-specific activation of T cells) could demonstrate their immunomodulatory capacity, however, further studies need to be performed to identify targeted genes in different immune cells. In a very recent study (81) equine amniotic MSCs were primed with a combination of IFNγ and TNFα demonstrating no additional immunosuppressive activity in an inflammatory *in vitro* model, compared to non-primed cells. Even though this study was performed with lower concentrations of IFNγ and TNFα on a different type of MSCs, it remarks that further research is necessary to confirm our insights.

In conclusion, our results have demonstrated that IFNγ and TNFα had a synergistic effect on IDO1 secretion, being a strong and efficient endMSCs priming strategy. Under this priming condition, surface molecules can be modified showing an increase of adhesion molecules, cytokine receptors and immune checkpoint receptors that may alter their biodistribution as well as their immunomodulatory activity. Here we hypothesize that IFNγ/TNFα may enhance the anti-inflammatory capacity of MSCs, as well as the migration/adhesion to inflammatory tissue. Finally, according to NGS and *in vitro* assays, IFNγ/TNFα-priming of endMSCs produce secretomes with a potent therapeutic/immunomodulatory potential.

## Data Availability Statement

The datasets generated for this study can be found in Sequence Read Archive (SRA) data with the accession number PRJNA664968^9^.

## Author Contributions

MÁP and JGC conceived and designed the experiments. MÁP, VÁ, EL, FM, MP, and JGC performed the experiments and analyzed the data. MÁP and JGC wrote the manuscript. All authors contributed to the article and approved the submitted version.

## List of abbreviations

BSA: Bovine Serum Albumin
CPM: Counts Per Million
DMEM: Dulbecco’s Modified Eagle’s medium
EDGE: Empirical analysis of Differential Gene Expression
ELISA: Enzyme-Linked ImmunoSorbent Assay
endMSCs: Endometrial-derived stromal/Mesenchymal Stem Cells
endMSCs*: IFNγ/TNFα-primed Endometrial-derived stromal/Mesenchymal Stem Cells
endMSCs: Endometrial-derived stromal/Mesenchymal Stem Cells
EVendMSCs: Extracellular Vesicles from Endometrial-derived Mesenchymal Stromal/Stem Cells
EVs: Extracellular Vesicles
FBS: Fetal Bovine Serum
FDR: False Discovery Rate
GO: Gene Ontology
IDO1: Indoleamine 2,3-dioxygenase
IFNγ: Interferon gamma
LPS: Lipopolysaccharide
MFI: Mean Fluorescence Intensity
FMO: Fluorescence Minus One
MSCs: Mesenchymal Stromal Cells
NGS: Next Generation Sequencing
PBLs: Peripheral blood lymphocytes
PBS: Phosphate buffered saline
PCA: Principal Component Analysis
Poly (I:C): Polyinosinic: polycytidylic acid
QPCR: Quantitative Real-Time Polymerase Chain Reaction
SD: Standard Deviation
SAR: Sequence Read Archive
S-endMSCs: Secretome from Endometrial-derived Mesenchymal Stromal/Stem Cells
S-endMSCs*: Secretome from IFNγ/TNFα-primed Endometrial-derived Mesenchymal Stromal/Stem Cells
SI: Stain Index
TLR: Toll-Like Receptor
TNFα: Tumor Necrosis Factor alpha
UMI: Unique Molecular Index.

## Acknowledgments

Lymphocyte activation assays were performed by ICTS Nanbiosis (Unit 14 at CCMIJU). The authors acknowledge the contribution of Joaquín González for the technical support in figures handling.

## 1. Figure legends

**Supplementary table 1. Panel of human monoclonal antibodies used for the phenotypic characterization of endMSCs and IFN**_γ_**/TNF**_α_**-primed endMSCs by flow cytometry**. The table includes relevant information of these markers.

**Supplementary table 2. The most abundant miRNAs in secretomes (S-endMSCs and S-endMSC*)**.

**Supplementary table 3. The top ten of miRNAs with the highest fold change in secretome samples**.

## Footnotes

1 http://geneontology.org/

2 https://reactome.org/

3 http://www.bioinformatics.babraham.ac.uk/projects/fastqc/

4 https://biit.cs.ut.ee/clustvis/

5 https://www.mirnet.ca/

6 http://www.lirmed.com/tam2/

7 https://ccb-web.cs.uni-saarland.de/mirtargetlink/

8 http://www.ncbi.nlm.nih.gov/bioproject/664968

9 http://www.ncbi.nlm.nih.gov/bioproject/664968

